# Phytofabrication of gold nanoparticles (AuNPs) via Wedelia chinensis Internodal extract: Characterization, Antioxidant, Antimicrobial, Hemolytic assay and Methylene Dye degradation properties

**DOI:** 10.1101/2025.06.17.660090

**Authors:** Brajesh Chandra Pandey, Ashish Gupta, Renu Kushwaha, Mohd Yaseen, Gopal Nath, Nishi Kumari

## Abstract

*Wedelia chinensis* is a medicinal herb of Asteraceae family. Preliminary phytochemical screening of internodal extract was done by using its aqueous, methanolic and ethanolic extracts. Preliminary phytochemical screening showed the presence of several pharmaceutically important compounds such as alkaloids, tannins, anthraquinons, saponins, flavonoids, terpenoids, phenols, etc. Aqueous internodal extract was used for the biosynthesis of gold nanoparticles. The characterization of gold nanoparticles were done by using a range of analytical techniques, such as energy-dispersive X-ray spectroscopy (EDS, EDX), X-ray diffraction (XRD), transmission electron microscopy (TEM), dynamic light scattering (DLS), Fourier transformed infrared spectroscopy (FTIR), scanning electron microscopy (SEM). The stability and other characteristics were ascertained using a UV-Vis spectrophotometer. The plasmon resonance occurred at the wavelength of 530 nm, thus showing the formation of gold nanoparticles. X-ray diffraction demonstrated the existence of Au-rich (fcc) phases in gold nanoparticles. FTIR analysis clearly indicated the participation of active constituents of internodal extract in the conversion of gold ions into AuNPs. DLS showed the size distribution (99.71 nm) of the suspended particles. There was variations in the sizes and shapes of the nanoparticles. The zeta potential of synthesized AuNPs was observed as −5.12 mV, thus showing the stability of the synthesized nanoparticles. In TEM study, uniform distribution of nanoparticles was seen with some agglomerates and average size of nanoparticles was observed in between 30-40 nm. The antioxidant potential of internodal extract was slightly higher than the synthesized gold nanoparticles. The green-synthesized AuNPs along with antibiotics demonstrated strong synergistic effect against several bacterial strains such as *Escherichia coli* and *Bacillus subtilis*. Gold nanoparticles showed good hemolytic activity against rat erythrocytes (90%). Gold nanoparticles also showed catalytic activity that reduces methylene blue at low concentration (1 mg).

**Highlights:**

➢ Internodal extracts showed the presence of several active constituents in preliminary phytochemical screening.
➢ Internodal extract was used for the synthesis of gold nanoparticles in one step process.
➢ Synergistic action of AuNPs and antibiotics were reported against many pathogenic bacterial strains.
➢ AuNPs showed its potential as antioxidant.
➢ Hemolytic effect of nanoparticles was reported on rat blood sample.
➢ Nanoparticles showed dye reducing activity thus enhancing its application in the purification of water bodies.

Fig.
Graphical abstract of AuNPs

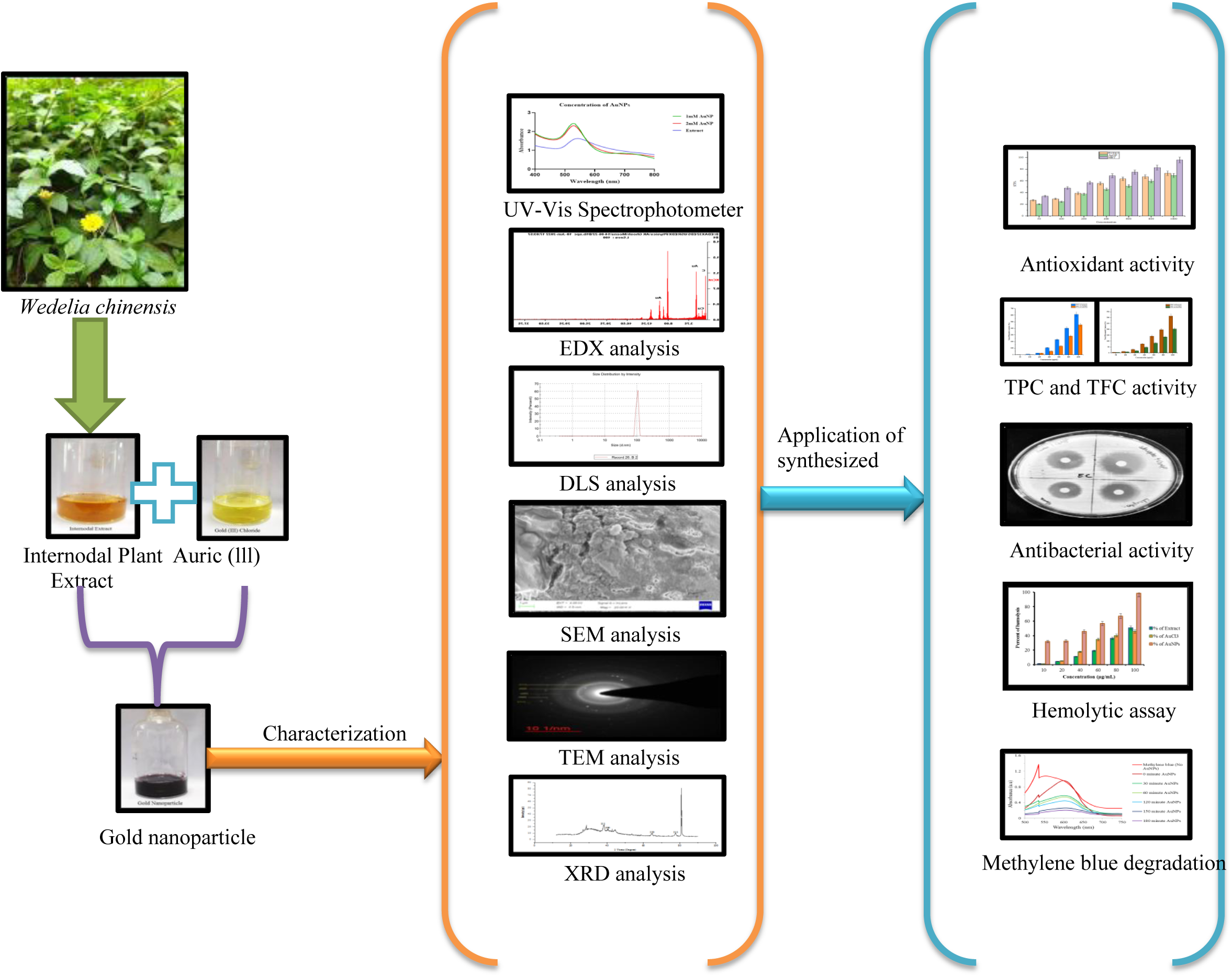

## 1. Introdution

The nanotechnology emerged as most promising area of science due to its large number of applications. Nanoparticles are broadly divided into different categories such as carbon-based NPs, metal NPs, ceramics NPs, semiconductor NPs, polymeric NPs, lipid-based NPs (**1**). Scientists prefer metal nanoparticles due to their enhanced features such as facet, shape and size controlled synthesis (**1**). Gold, silver, platinum and copper nanoparticles are some commonly used metal nanoparticles. They display wider applications due to their improved optical, opto-electrical, catalytic characteristics with significant biological activities (**2**).) Wider applications of gold nanoparticles in medical and non-medical fields are due to their many unique features such as inertness, biocompatibility and low toxicity. Some of its non-medical applications are removal of bad smell, removal of harmful carbon monoxide from rooms, emission management, water purification, power cell, etc. It has wider applications in health sector. Due to their small size, the particle can enter tissues and attack immune cells, thus making it suitable for immunotherapy purpose (**3**). Gold nanoparticles are suitable in different types of cancer therapies such as chemo-radio-therapy, thermo-chemo-radio – therapy, gene therapy. AuNPs show multifunctional ability and are widely used in biomolecular screening, cancer therapy, photo thermal therapy, labelling of proteins, gene and DNA delivery (**4–5**). Morphology of gold nanoparticles can be modified by altering the reaction conditions and thus diverse shapes and sizes of NPs can be obtained. For the synthesis of nanoparticles, researchers are generally preferring biological extracts. Biological methods are less complicated, eco-friendly, cost-effective and more productive (**6**). Metabolites present in plant extracts cause bio-reduction of metal ions to nanoparticles. These metabolites also provide stability to the nanoparticles.

*Wedelia chinensis* is a medicinal herb of Asteraceae family. Its different parts are being used for the treatment of cancer, ulcer, inflammation, wound healing, CNS disorders, etc. as traditional medicine in Ayurveda, Unani and Sidhha systems (**7**). *W. chinensis* shows significant presence of active constituents such as flavonoids, diterpenes, triterpenes, saponins, etc (**8**). The synthesis of silver nanoparticles of *W. chinensis* have been reported from leaf (**7**) and flower (**9**) extracts. There is an utmost need to synthesize gold nanoparticles and to explore its applications. In the present study, internodal extracts have been taken for the synthesis of AuNPs. Preliminary phytochemical screening of the internodal extracts was also done to study its medicinal potential. The characterization of gold nanoparticles was done by different assays and their biological activities were assessed.

## 2. Experimental

### 2.1. Chemicals and Reagents

Gold (III) chloride trihydrate (HAuCl_4_. 3H_2_O) was obtained from Hi-Media Chemical Reagent Co. Ltd. Molecular grade ethanol, sodium chloride (NaCl), hydrochloric acid (HCl), ascorbic acid and sodium hydroxide (NaOH) were purchased from Hi-media. Chemicals for antioxidant and antimicrobial activity including 2,2-diphenyl-1-picrylhydrazyl (DPPH) and antibiotics (streptomycin, Gentamycin, Kanamycin and Streptomycin) were obtained from Hi-Media and SRL, respectively. The chemicals used for total phenol content (TPC) and total flavonoid content (TFC): Rutin (standard) and Gallic acid (Standard) was purchased from Hi-Media. Folin-Ciocalteu’s phenol reagent was purchased from Sigma-Aldrich, sodium carbonate, aluminium chloride (AlCl_3_), sodium nitrite (NaNO_2_) was obtained from Hi-Media. Muller Hinton Broth (MHB) and Muller Hinton Agar (MHA) were purchased from companies including SRL and Hi-Media.

### 2.2. Extract Preparation

Fresh internodes of *Wedelia chinensis* (**Fig. 1**) (**Specimen no. Astera. 2024/01: accession number was provided by Prof N K Dubey, Department of Botany, BHU**) were obtained from the Ayurvedic garden of Dravyaguna Department, IMS, BHU. 100 gm of internodes were thoroughly washed with tap water and then rinsed with double distilled water. After washing, plant materials were shade dried for about one week and then crushed to obtain fine powder. Powdered plant extracts were used both for phytochemical screening and gold nanoparticle synthesis.

**Fig. 1.**
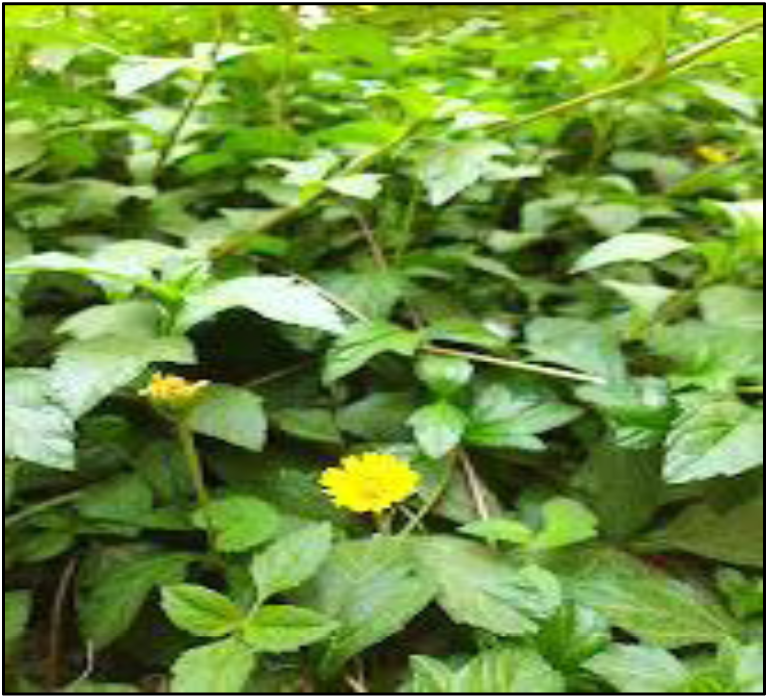
*Wedelia chinensis* plant along with flower and internode.

### 2.3. Preliminary phytochemical screening

#### 2.3.1. Plant Extract Preparation

Internodal part of the *Wedelia chinensis* were dried and was made as powder.

##### 2.3.1.2. Preparation of Extract

10 gm of dried powder of *W. chinensis* was suspended in 100 mL of water, ethanol and methanol solvents, respectively. Extraction was done using Soxhlet apparatus for 3 hrs at a specific temperature for each solvent. Further, the extract obtained after the mentioned process was preserved in the refrigerator in glass bottle throughout the experiment (i.e. for qualitative analysis). Qualitative phytochemical screening of internodal extracts of *Wedelia chinensis* was performed by using protocols (**10**). The methods employed for each phytochemical test are summarized below.

#### 2.3.2. Alkaloids

Mayer’s test was performed by adding a few drops of Mayer’s reagent to the extracts prepared in different solvents. The formation of green color or white precipitate indicated the presence of alkaloids.

#### 2.3.3. Tannins

The presence of tannins was determined by using the Ferric chloride (5%) test. Few drops of ferric chloride solution were added to different extract solutions. The formation of blue black or greenish black color is an indication of tannin presence.

#### 2.3.4. Anthraquinones

Brontrager’s test was conducted by adding dilute hydrochloric acid in each extract solutions (5 mL) of plant extract followed by Benzene and then 10% of Ammonia (1 part of ammonia, add 9 part of water). The appearance of pink or red color precipitate shows the presence of anthraquinones.

#### 2.3.5. Reducing sugars

Benedict’s test was performed to indicate the presence of reducing sugar. In this process Benedict’s reagent was added to 5 mL each extract solutions and then it was heated. Formation of a brick red precipitate indicates the presence of reducing sugars.

#### 2.3.6. Saponins

Froth test was performed by shaking extract solution vigorously for 10 to15 minutes. 3 mL of extract solutions was added with distilled water and shaken vigorously. The formation of a thick foam layer indicates the presence of saponins.

#### 2.3.7. Flavonoids

Alkaline reagent test was performed by adding 2N of NaOH to 3 mL of each extract solutions. Formation of a yellow color indicates the presence of flavonoids.

#### 2.3.8. Phlobatannins

Phlobatannins was determined by using dilute HCl. Plant extract solution of each solvent was treated with 2% of hydrochloric acid. Formation of a red color precipitate indicates the presence of phlobatannins.

#### 2.3.9. Steroids

Liebermann–Burchard test was conducted by adding chloroform and few drops of concentrated sulfuric acid to 4 mL of different extract solutions. The formation of brown ring indicates the presence of steroids and bluish brown ring indicates the phytosteroids.

#### 2.3.10. Terpenoids

The test was performed by adding chloroform and concentrated sulfuric acid to 2 mL of solvents containing plant extracts. Formation of a red brown color at the interface indicates the presence of terpenoids.

#### 2.3.11. Coumarins

The NaOH test was performed by treating 5 mL of each extract sample with 10% of sodium hydroxide solution. Formation of a yellow color indicates the presence of coumarins.

#### 2.3.12. Protein and amino acids

Ninhydrin test was conducted by adding 0.2% Ninhydrin to 5 mL of each extract sample and the samples were heated for 10 min. Formation of blue color indicates the presence of protein and amino acids.

#### 2.3.13. Anthocyanin

The presence of anthocyanin was determined using NaOH. 5 mL of each extract was treated with 2N NaOH and was heated for five minute at 100⁰C. Formation of bluish green color shows the presence of anthocyanin.

#### 2.3.14. Carbohydrates

Molisch’s test was performed for the determination of Carbohydrate. In this experiment, Molisch’s reagent was added to each extract solution followed by few drops of concentrated H_2_SO_4_. Formation of a purple or red ring at the interface determines the presence of carbohydrates.

#### 2.3.15. Quinines

The presence of Quinines was confirmed by adding sulphuric acid to extract solution. The formation of red color indicates the presence of Quinines in the plant sample.

#### 2.3.16. Phenols

Ferric chloride test was performed for the test of Phenols. Few drops of 10% of FeCl_3_ was added to each extract solution (final volume is 5 mL). Formation of blue or green color indicates the presence of phenols.

#### 2.3.17. Cardiac Glycosides

Keller Killiani test was performed for Cardiac glycosides. Few drops of 1 % of FeCl_3_ solution and 5% of glacial acetic acid were added to 5 mL of each extract. This solution was under layered with concentrated sulphuric acid. Formation of brown ring at the interface indicates he presence of Cardiac glycosides.

### 2.4. Synthesis of Gold nanoparticles

Fine powder (10 g) of *W. chinensis* was dissolved in 100 mL of double distilled water in a beaker and was stirred on a magnetic stirrer (200 rpm) for overnight. After the incubation period, extract was filtered with the help of a muslin cloth and was dried in rotary evaporator to make the extract moisture free. The dried extract powder (100 mg) was dissolved in 1 mL of double distilled water. Different concentrations (0.1, 0.2 or 0.4 M) of chloroauric acid (volume 10 mL) were mixed with 1 mL of extract. The change of color from light yellow to reddish-brown was observed after incubation period (**Fig. 2**). To optimize synthesis conditions, different temperature (like 25^0^C, 50^0^C and 100^0^C), pH (2, 7 and 12) and reaction time (2 hrs, 6 hrs and 12 hrs) were studied and suitable condition was observed under spectroscopy. To get suitable concentration of AuNPs for characterization studies, different concentrations of AuNPs (1 mM and 2 mM) were taken and compared with extract solution. Optimized conditions of synthesized NPs were used for further characterization and biological activity studies.

**Fig. 2.**
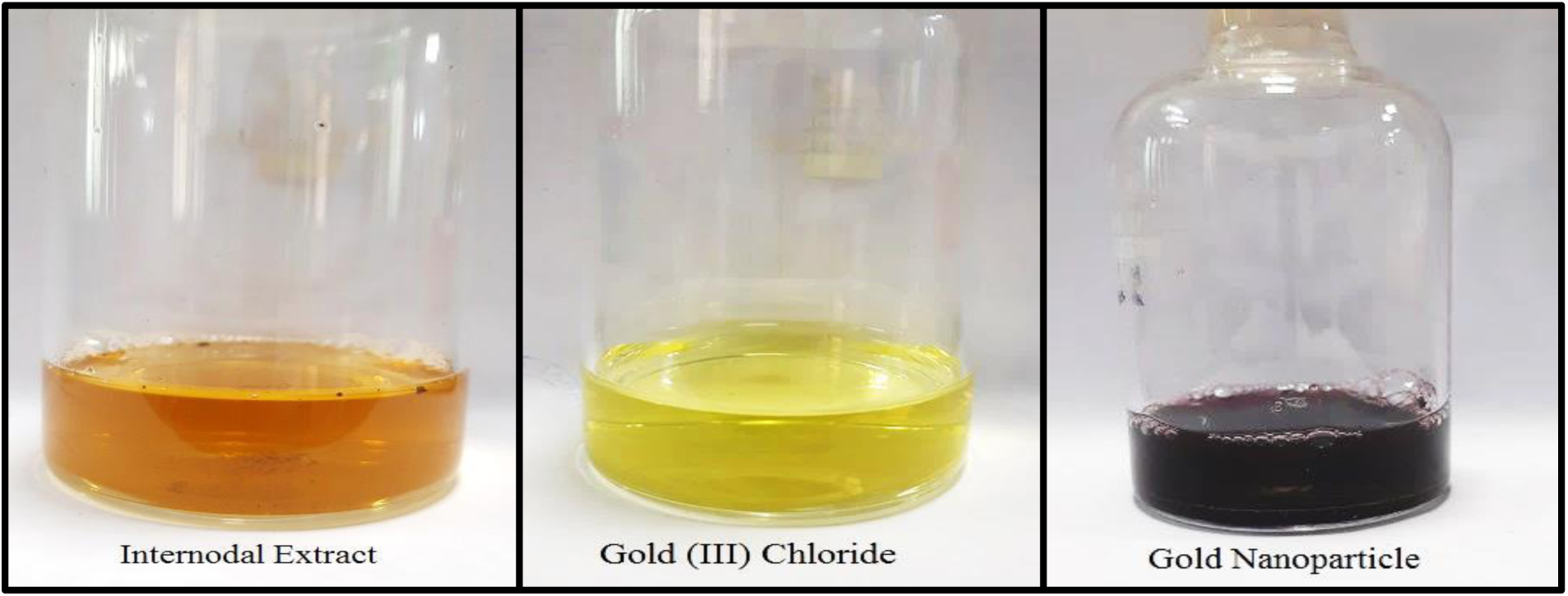
Synthesis of Gold nanoparticle from the Internodal extract of *Wedelia chinensis*.

### 2.5. Characterization of synthesized Gold nanoparticles

#### 2.5.1. Instruments used

UV–Visible spectroscopy (UV-2600i UV-2700i Shimadzu Corporation), Dynamic light scattering (Malvern Instruments Ltd. for Zeta Sizer), Transmission Electron microscope TECHNAI (JEM-2100F, JEOL), Energy-dispersive X-ray analysis (EDX), Scanning electron microscope (GemniSEM 460), X-ray diffraction analysis (XRD-6000; Shimadzu, Kyoto, Japan), Fourier transformed infrared spectroscopy (JASCO FT/IR-6600 machine). For characterization of nanoparticles, the methodology was same as followed by the protocols (**9**).

##### 2.5.1.1. UV–Visible spectroscopy

UV–Visible spectrometric analysis is used for nanoparticle characterization. The observed absorption peaks in the UV–Visible spectra provide crucial insights into nanoparticle synthesis and stability. UV– Visible spectroscopy in the range of 400–800 nm was used as the confirmatory technique for the synthesis of Gold nanoparticles. The samples were placed in quartz cuvette. The UV–Visible spectrum was recorded for *W. chinensis* internodal aqueous extract and AuNPs. Different parameters of NP synthesis were analyzed such as temperature (like 25^0^C, 50^0^C and 100^0^C), pH (2, 7 and 12), stability with reaction time (2 hrs, 6 hrs and 12 hrs). Optimum concentration of AuNPs were also checked.

##### 2.5.1.2. Dynamic light scattering

DLS is used to measure the size distribution of gold nanoparticles. DLS can provide information about the diameter, size distribution, and the stability of nanoparticles in a solution. The technique can be used to determine the size of the nanoparticles, including both the core and the surface coating in the suspension.

##### 2.5.1.3. Zeta potential

Zeta potential analysis measures the surface charge of the gold nanoparticles. Zeta potential analysis can provide information about the stability of nanoparticles in solution, as nanoparticles with higher zeta potentials are more stable against aggregation and precipitation (**11**).

##### 2.5.1.4. TEM analysis

Transmission Electron Microscopy (TEM) is used to determine microstructure of materials at high resolution. TEM images of AuNPs can reveal important information about their morphology, size, and distribution. This information can be used to determine the properties of the nanoparticles and their potential applications.

##### 2.5.1.5. SEM analysis

Scanning Electron Microscopy (SEM) is used to determine the morphology and size of the formed gold nanoparticles. The powder form of synthesized AuNPs was used to obtain SEM images and to determine the shape and size of nanoparticles. The experiment was conducted with a voltage of 4.0 kV (**12**).

##### 2.5.1.6. X-ray diffraction analysis

XRD is a technique used to study the structure of nanoparticles. XRD can provide important information about the size, shape, and crystalline structure of gold nanoparticles (**13**).

##### 2.5.1.7. FTIR spectroscopy

FTIR was used to identify the functional group composition of internodal aqueous extract of *W. chinensis* and AuNPs. The spectra of the internodal extract of *W. chinensis* and the synthesized AuNPs were compared with the data provided in the research articles to confirm the synthesis of AuNPs (**14**).

##### 2.5.1.8. EDX analysis

Energy-dispersive X-ray analysis (EDX) was used to identify the elemental composition of the Gold nanoparticles. An EDX study was performed in different regions of the synthesized AuNPs, which depicts the elemental composition of the gold nanoparticles (**15**).

##### 2.5.1.9. EDX Mapping

EDS Mapping was done to understand the spatial distribution of elements in a solid material. EDX is the combination of spatial resolution of electron microscope with the power of x-ray spectroscopy.

### 2.6. Antioxidant activity of the synthesized Gold nanoparticles

#### 2.6.1. DPPH activity

Methanol solution of DPPH (0.2 mM) discoloration test was performed to evaluate the radical scavenging capability of internodal extract and AuNPs and particularly, this discoloration technique was operated by DPPH reduction. The DPPH solution interacts with the different concentrations of extract, AuNPs and positive control (from 50 to 1000 μg/ml) as a reducing agent that the DPPH color changed from light yellow to reddish brown (the degree of color change shows the antioxidant capacity of the sample), after incubating the solution for 30 min in dark. The reading was taken at 517 nm to determine the change of color. Butylated hydroxytoluene (BHT) was taken as a positive control and DPPH radical scavenging activity was taken as an inhibition percentage (**16**). The experiment was performed in triplicate and the scavenge half of the DPPH free radical (IC_50_) was obtained, the inhibition graph was plotted against concentration.

#### 2.6.2. Estimation of TPC and TFC Folin-Ciocalteu’s Reagent Assay and AlCl_3_ Colorimetric Method

The total phenolic content of internodal plant extract and AuNPs were determined using the Folin – Ciocalteu method (**17**). Sample solutions (1 mg/mL) at varying concentrations (50, 100, 200, 400, 600, 800 and 1000 µg/mL) and freshly prepared 0.2 N Folin-Ciocalteu reagent (2 mL) were taken in the test. Then, 2 mL of the 75 g/L sodium carbonate (Na_2_CO_3_) was added into the test tubes and incubated for 30 minutes at room temperature. The absorbance of the reaction mixture was measured at 765 nm. Phenolic content was measured by using rutin calibration curve and was expressed as rutin equivalents (µg RE).

Total flavonoid content was measured by the method (**18**) with slight modification. Aluminum chloride solution prepared in 2% methanol was mixed with different concentrations (50, 100, 200, 400, 600, 800 and 1000 µg/mL) of experimental samples (Stock solution of internodal extract and AuNPs was taken as 1 mg/mL). Absorbance was measured at 415 nm, and the total flavonoid content was determined using a standard curve of Gallic acid and was expressed as Gallic acid equivalents (µg GAE).

### 2.7. Synergistic effect of Gold nanoparticles on different Bacteria strains

Antibacterial activities of green-synthesized gold nanoparticle and their combinations with different antibiotics (Rifampicin, Gentamycin, Kanamycin, and Streptomycin) were observed by the disc diffusion method (**19**). Six test organisms were taken for the experiment: *Klebisiella pneumoniae*, DH5α, MRSA (Methicillin-resistant Staphylococcus aureus), *Escherichia coli*, *Pseudomonas aeruginosa*, *Shigella boydii*. Mueller Hinton agar medium was prepared and autoclaved. About 25 mL media were poured into sterile petri plates and the plates were allowed to solidify for 15 min. After that 10 µL of bacterial inoculum was plated uniformly on the petri plate surface and was allowed to dry for 10 min. Different combinations of extract, AuNPs and antibiotics was loaded on 2.5 mm diameter of sterile disc, which were placed on the surface of the medium and the plates were kept for incubation at 37⁰C for 24 hrs and the zone of inhibition (in mm) was calculated.

### 2.8. Hemolytic assay of Internodal extract and gold nanoparticle

#### 2.8.1. Erythrocytes suspension preparation

About 4-5 ml of blood was collected from a rat (Charles Foster Rat) in centrifuge tube having EDTA (Ethylenediamine tetra acetic acid). Blood was then centrifuged at 5000 rpm for 10 min. Supernatant (Plasma) was then discarded and the pellet containing RBCs was washed for 2-3 times by centrifugation at 5000 rpm for 5 minutes with phosphate buffer saline (PBS) having pH 7.2. The pellet was then kept in 0.5% normal saline.

#### 2.8.2. Cytotoxicity study by Hemolytic activity

Cytotoxic study of hemolytic activity was performed with the help of spectrophotometer (**20**). Internodal extract of the plant and AuNPs were taken as a stock solution (1mg/ml each) in phosphate buffer. Working solution of different concentrations (10, 20, 40, 60, 80 and 100 μg/ml) of plant extract and AuNPs were prepared from the stock solution. 1 ml cell suspension was added in different concentrations of plant extracts and gold nanoparticles (volume of plant extract and gold nanoparticles were 1ml). The whole reaction mixture was incubated at 37°C for 30 min. The whole reaction mixture was undergone for centrifugation at 2000 rpm for 5 min in a centrifuge. Absorbance of supernatant containing free hemoglobin (hemoglobin present outside of RBCs and also formed as a liquid part of the blood) was observed in a UV-Vis spectrometer at a wavelength of 540 nm. Triton X-100 (0.1%) was taken as a positive control and phosphate buffer saline (PBS) was used as the negative control. The percentage of hemolysis was calculated by the formula;

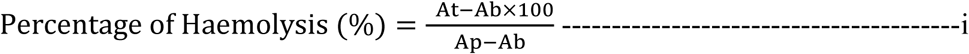

Where,

At is absorbance of test sample,

Ap is absorbance of the positive control,

Ab is absorbance of negative control.

### 2.9. Reduction of Methylene Blue Dye

A stock solution of methylene blue (0.05 mg/mL) was prepared by dissolving 5 mg of the dye in 100 mL of deionized water. The experiment was conducted by mixing 1mL of freshly prepared aqueous sodium borohydride (NaBH_4_) solution (0.1 M), 30 mL of the methylene blue stock solution, and 30 μL of AuNPs (1 mg/mL). A mixture without nanoparticles was used as a negative control. Catalytic degradation of the dye was assessed by withdrawing 3 mL aliquots at different time intervals and monitoring the dye’s characteristic absorption band at 625 nm using UV-visible spectrophotometer. The catalytic efficiency was measured according to the following equation:

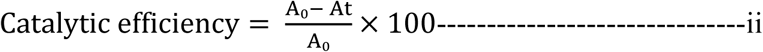

Where,

Ao is the absorbance of the Methylene blue dye solution (before AuNPs addition),

At is the absorbance of the Methylene blue dye solution at time t in the presence of gold nanoparticles.

### 2.10. Statistical analysis

The results of this investigation were presented as the mean ± standard of the mean (SEM) or SD. A statistical analysis was performed using data analyses function using one-way analysis of variance (ANOVA). The graphs were drawn in Origin Pro. All data were statistically analyzed using the GraphPad Prism software, version 8.0.1 (GraphPad Software Inc.).

## 3. Results and discussion

### 3.1. Phytochemical analysis of Internodal extracts of *Wedelia chinensis*

Preliminary phytochemical screening of internodal extracts of *W. chinensis* shows the presence of alkaloids, tannins, anthraquinones, saponines, steroids, phlobatannins, terpenoids, protein and amino acids, anthocyanin, phenols and coumarins (**Table. 1**). Most of these phytochemicals are of therapeutic values showing antioxidant, antimicrobial, immunomodulatory, antidiabetic and anticancer activities (**23**).

**Table. 1.** Different Phytochemical test for Wedelia chinensis Internodal plant extract.

|  |  |  |  |  |
| --- | --- | --- | --- | --- |
| Phytochemical | Test Performed | Result |  |  |
|  |  | Methanolic extract | Aqueous extract | Ethanollic extract |
| Alkaloids | Mayer's test | + | + | + |
| Tannins | Ferric chloride test | + | + | – |
| Anthraquinones | Brontrager test | + | + | – |
| Reducing sugars | Benedict's test | – | – | – |
| Saponins | Frothing test | + | + | – |
| Flavonoids | Alkaline reagent test | – | + | – |
| Phlobatannins | HCl test | + | + | + |
| Steroids | Liebermann–Burchard test | + | + | – |
| Terpenoids | Salkowski test | + | + | – |
| Coumarins | NaOH test | + | + | + |
| Protein and amino acids | Ninhydrin test | + | + | – |
| Anthocyanin | NaOH test | + | + | – |
| Carbohydrates | Molisch and Fehling's tests | – | + | – |
| Quinines | H<sub>2</sub>SO<sub>4</sub> test | – | – | + |
| Phenols | FeCl<sub>3</sub> test | + | + | + |
| Cardiac Glycosides | Killer killiani test | – | + | – |

### 3.2. Synthesis of NPs

Synthesis of Gold Nanoparticles from aqueous extract of *W. chinensis* was visually confirmed by the change in color of reaction mixture from light yellow to reddish brown. The change in color of the reaction mixture from pale yellow to ruby red color is the confirmation of gold nanoparticle synthesis (**24**). Green biosynthesis of AuNPs has several advantages-easy availability of plant materials, cost-effective, non-toxic, eco-friendly and synthesis at room temperature. In present study, internodal extract of *W. chinensis* is rich reservoir of high value phytochemicals. The phytochemicals present in the crude plant extracts serve dual purposes as reducing as well as stabilizing agents (**25–26**).

### 3.3. Characterization

#### 3.3.1. UV–Visible spectrometric analysis

UV–Visible spectra show plasmon resonance of AuNPs between 500-600 nm, thus confirming the presence of gold nanoparticles (**Fig. 3**). Several researchers observed peak formation by gold nanoparticles in same range under spectroscopic study (**27–28**). In present work, high temperature (100° C) was observed as suitable condition for NP synthesis. It was reported increase in the peak of the absorption spectra of nanoparticles with rise in temperature (**Fig. 3A**) (**29**). Reaction time plays a key role in the biosynthesis of nanoparticles (**30**). Optimum reaction time for the synthesis was 12 hrs (**Fig. 3B**). We also check the effect of pH on the formed nanoparticles and we observed that at alkaline pH the absorbance peak was highest and as we move to the acidic pH the absorbance starts decreasing and was found lowest at pH 2 (**Fig. 3C**). In contrast to our findings, several researchers observed acidic pH favorable for AuNP synthesis (**31–32**). 1 mM AuNPs showed most prominent peak in spectrometric study, when compared with 2 mM AuNP and the extract (**Fig. 3D**). The effects of Ultraviolet light duration on the AuNPs were also studied for different time intervals and it shows maximum absorbance at 30 minute, but the absorbance decreases for longer exposure of UV (**Fig. 3E**).

**Fig. 3.**
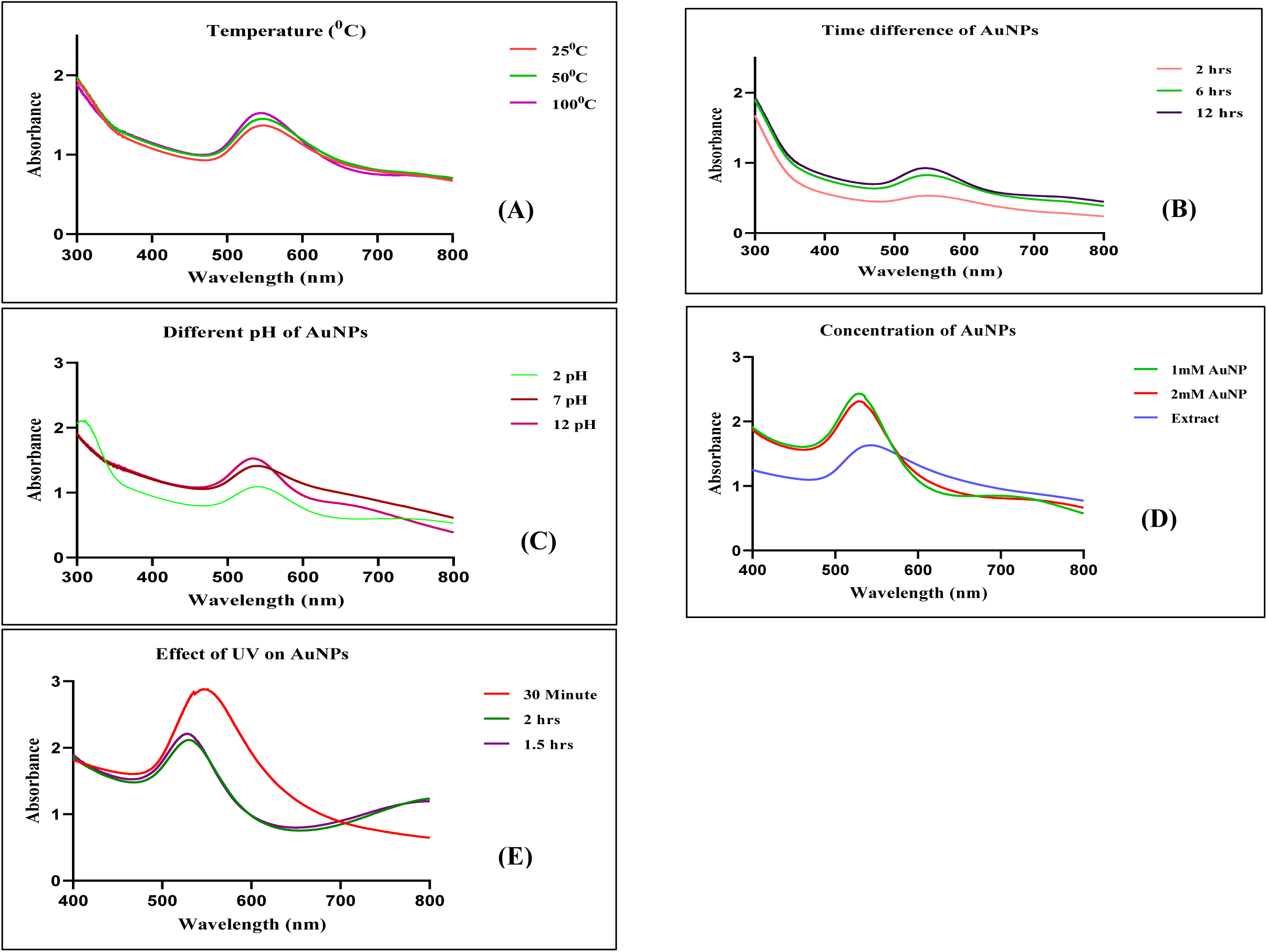
(**A**) Formation of AuNPs at different temperatures, **(B)** Formation of AuNPs at different time intervals, **(C)** Effect of different pH on the formation of AuNPs, **(D)** Effect of different concentration on the formation of AuNPs and comparison with extract, **(E)** Effect of UV on the stability of AuNPs.

#### 3.3.2. FT-IR analysis

FTIR was done to confirm the functional groups responsible for the reduction of the Au ions. A strong peak was observed at 3410 cm^−1^, indicating the presence of intramolecularly bonded alcohol groups (**33**). Similarly, the sharp peak at 1629 cm^−1^ indicates unsaturation in the plant extract. The peaks between 600 cm^−1^ and 500 cm^−1^ responsible for the carbon halogen bonds (**Table. 2**). Peaks such as 2931, 2344, 1655, 1032, etc. represents Carboxylic acid, Carbon di-oxide, Alkene, Sulfoxide, etc. The functional groups participated in the reduction and stabilization of Au^+^ to form AuNPs. The FTIR spectra of the internodal plant extract and AuNPs are shown in **Fig. 4**.

**Fig. 4.**
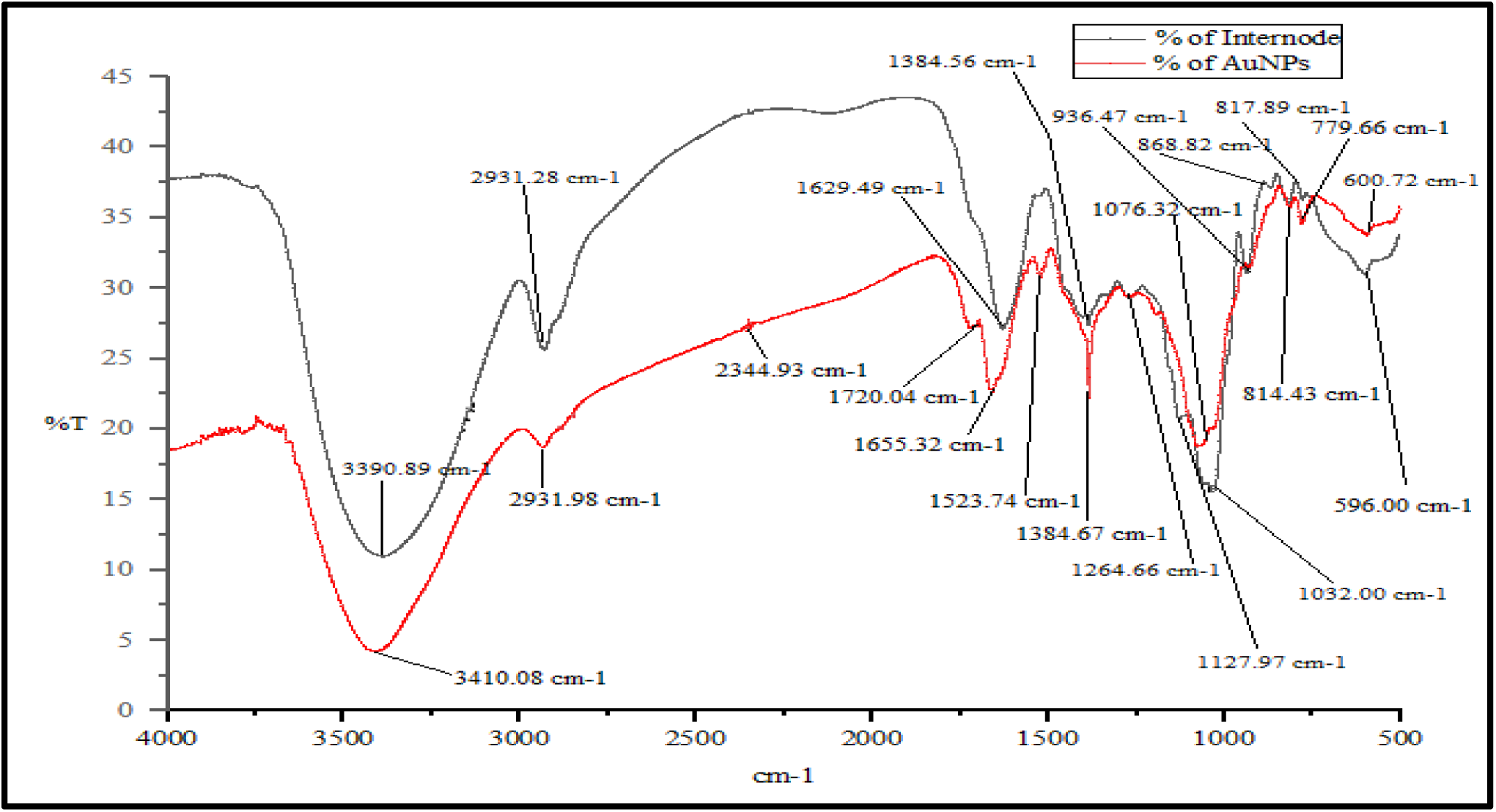
FTIR spectra of Internodal plant extract and AuNPs.

**Table. 2.** Different peak positions with their functional groups of AuNPs, and Internodal Extract of W. chinensis.

|  |  |  |  |  |
| --- | --- | --- | --- | --- |
| Biosynthesized AuNPs of *Wedelia chinensis* Internode |  |  | *Wedelia chinensis* Internodal Extract |  |
| Peak | X ($\text{cm}^{-1}$) | Functional Group | X ($\text{cm}^{-1}$) | Functional Group |
| 1. | 3410.08 | O-H stretching (Alcohol) | 3390.89 | O-H stretching (Alcohol) |
| 2. | 2931.98 | O-H stretching (Carboxylic acid) | 2931.28 | N-H stretching (Amine salt) |
| 3. | 2344.93 | O=C=O stretching ( Carbon di oxide) | 1629.49 | C=C stretching (Alkene) |
| 4. | 1720.04 | C=O stretching (Aldehyde) | 1384.56 | S=O stretching (Sulfate) |
| 5. | 1655.32 | C=C stretching (Alkene) | 1127.97 | C-O stretching ( Tertiary alcohol) |
| 6. | 1523.74 | N-O stretching (Nitro compound) | 1032.00 | S=O stretching(Sulfoxide) |
| 7. | 1384.67 | S=O stretching (Sulfate group) | 936.47 | O-H bending (Ccarboxylic group) |
| 8. | 1264.66 | C-O stretching (Aromatic ester) | 868.82 | C-H bending (Trisubstituted) |
| 9. | 1076.32 | C-O stretching (Primary alcohol) | 817.89 | C=C bending ( Alkene) |
| 10. | 814.43 | C=C bending (alkene) | 596.00 | C-I stretching (Halo compound) |
| 11. | 779.66 | C-Cl stretching (Halo compound) |  |  |
| 12. | 600.72 | C-I stretching (Halo compound) |  |  |

#### 3.3.3. Dynamic light scattering and Zeta potential

**Fig. 5A** representing the particle size distribution of AuNPs obtained from the Dynamic light scattering (DLS) technique showing that the average size of AuNPs is 99.71 nm. Zeta potential provides the information about surface charge potential index along with the stability of the nanoparticles in the aqueous suspension (**34**). The zeta potential value of AuNPs was found to be – 5.12 mV (**Fig. 5B**). Negative value of zeta potential for nanoparticles indicates their stability. More negative value of Zeta potential shows more interparticle repulsion and stability of nanoparticles (**35**), whereas less negative charge shows more capability of nanoparticles in cell penetration (**36**).

**Fig. 5.**
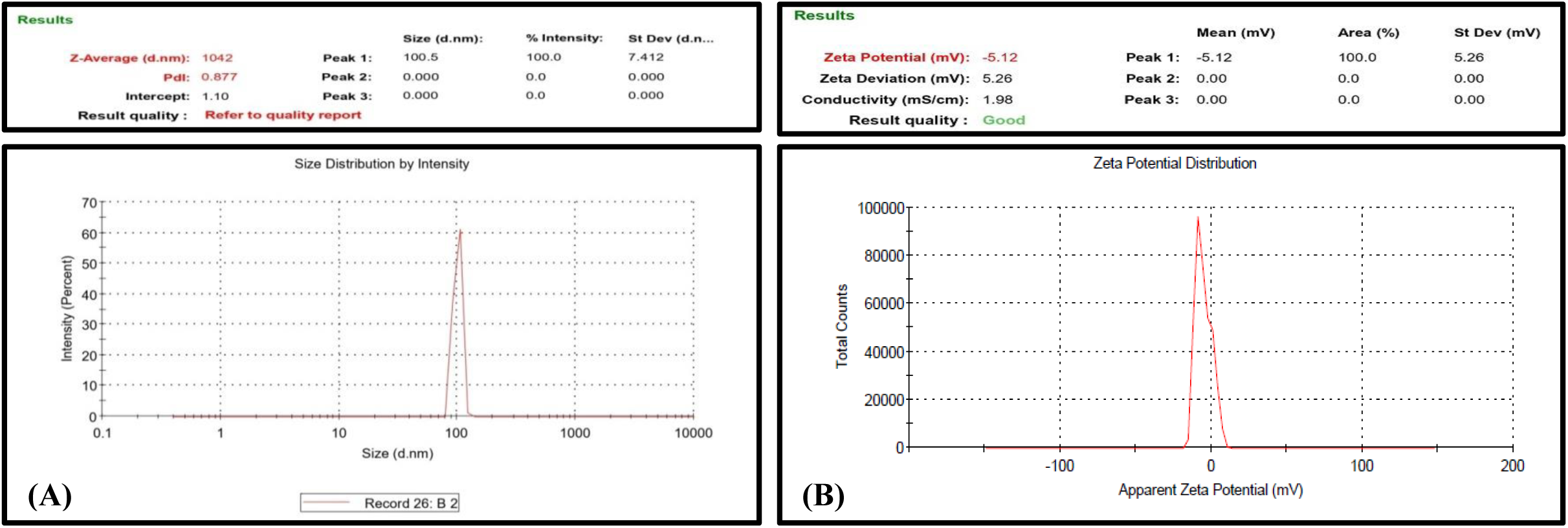
**(a)** Dynamic light scattering of AuNPs, **(b)** Zeta potential curve of AuNPs.

#### 3.3.4. EDX analysis

EDX provides the elemental composition as well as surface atomic distribution (SAD) of gold nanoparticles (**37**). The EDX analysis confirmed the presence of AuNP with a weight% of 63 and atomic% of 10.50 (**Fig. 6**). The presence of other elements (carbon, oxygen and copper) was also observed.

**Fig. 6.**
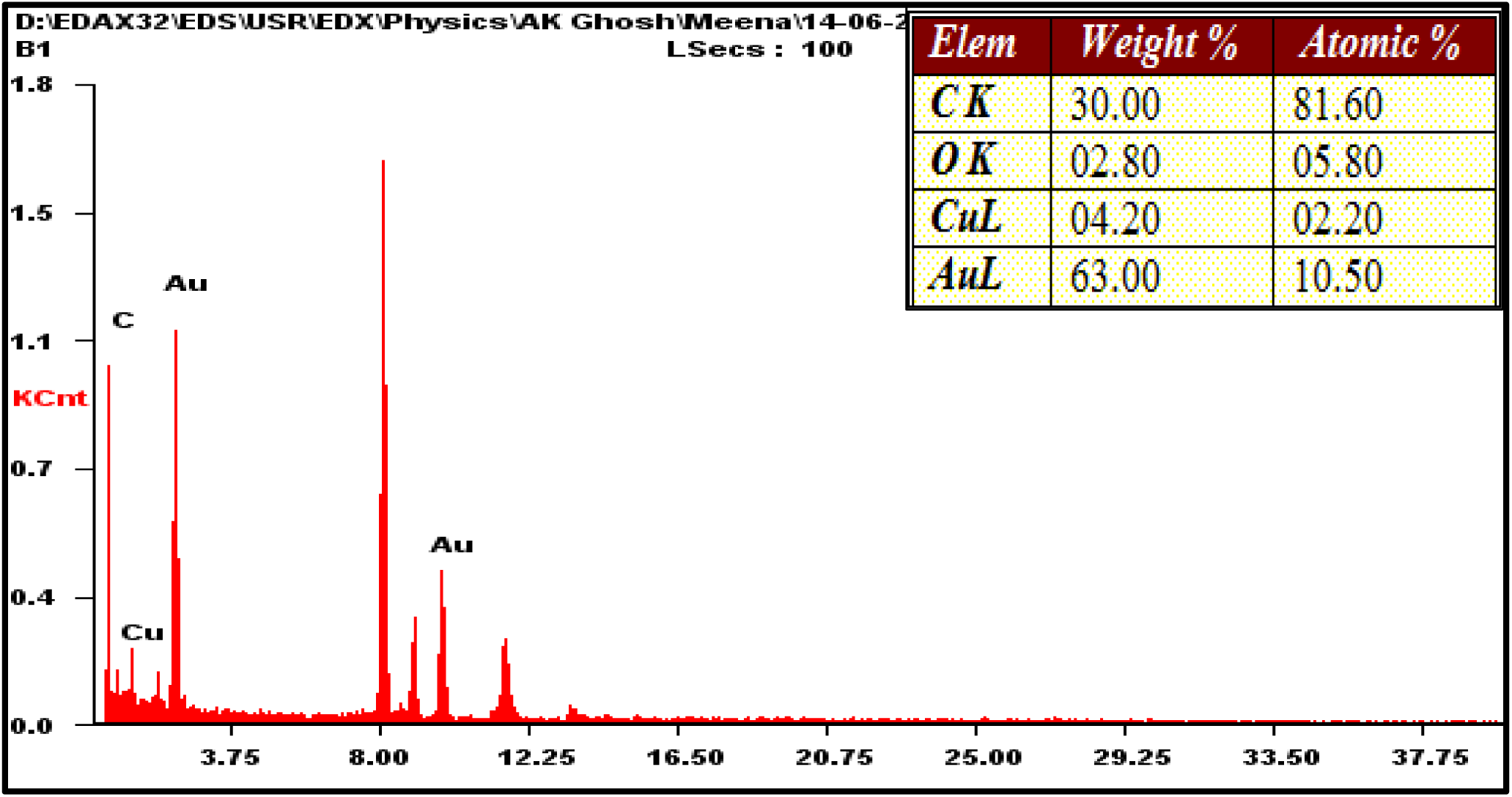
EDX spectrum of gold nanoparticles showed characteristic peaks.

#### 3.3.5. EDS mapping

EDS mapping reveals that the percentage of gold atom in the solution was 62.78%. Other elements were also present such as Carbon (10%), Oxygen (15%), Nitrogen (2.22%), Sodium (4%) and Potassium (6%). EDS mapping showed uniform distribution of gold atom on the internodal extract of *Wedelia chinensis* (**Fig. 7**).

**Fig. 7.**
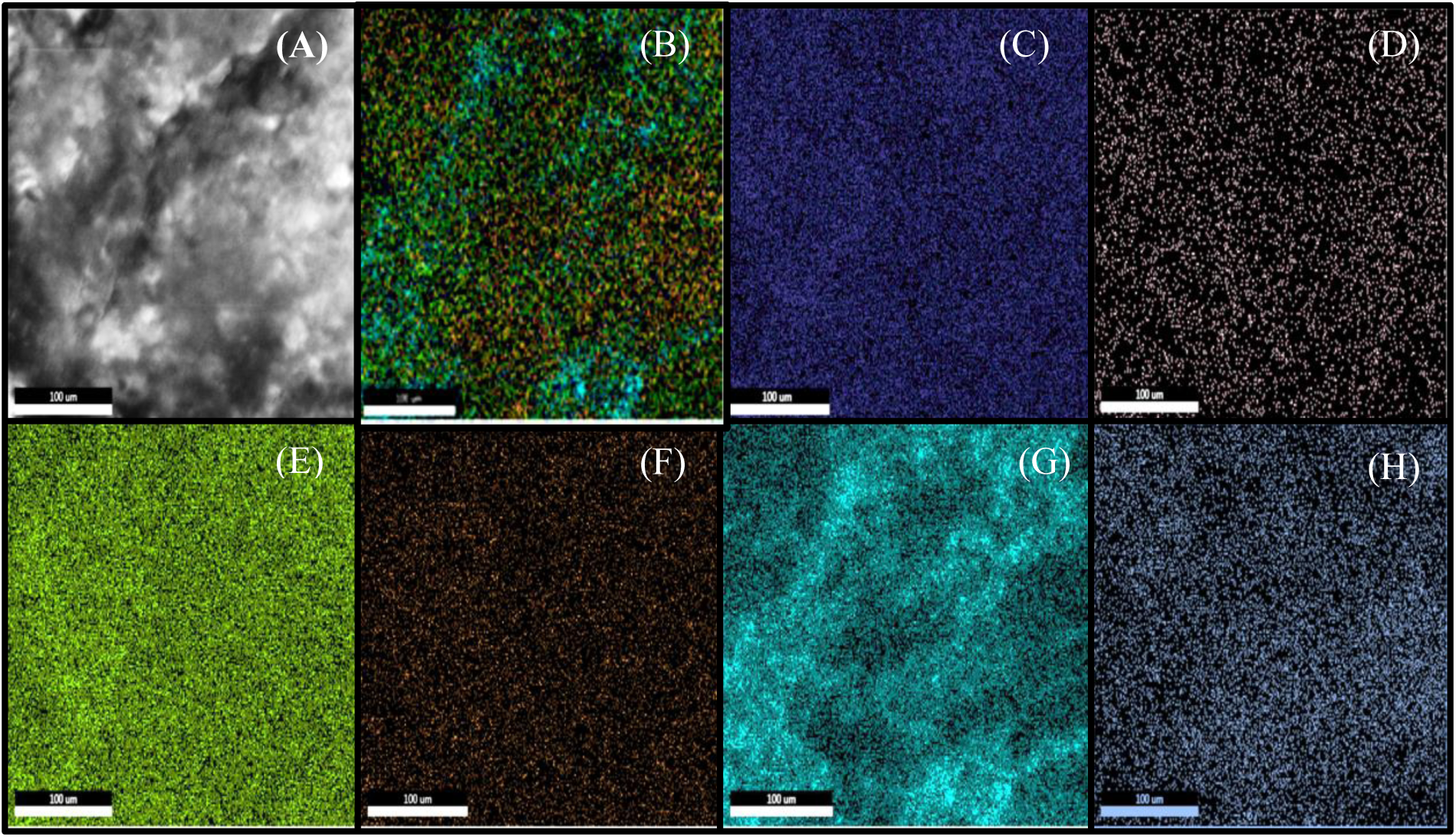
Showing EDS map of spectral images of Gold chloride loaded on the internodal extract of *W. chinensis*. (A) EDS image of treated internodal extract with gold chloride (Au (lll) Cl); (B) distribution of element overlay; (C) distribution of carbon element; (D) distribution of nitrogen element; (E) distribution of oxygen element; (F) distribution of sodium element; (G) distribution of potassium element; (H) distribution of gold element.

#### 3.3.6. TEM

TEM is used for analysis and size distribution histogram (the (200 nm, 100 nm, 50 nm, 20 nm, 0.5 nm, 10 1/nm scale). The study showed the presence of spherical, triangle, Pentagonal and Hexagonal morphologies of the formed nanoparticles with an average size of distribution of particle were in between 30-40 nm (**Fig. 8**). The size and stability of the formed AuNPs were stable and not agglomerated. The result obtained from DLS on comparing with the result of TEM is not consistent. The average size difference between DLS and TEM has been studied in previous research (**38–39**).

**Fig. 8.**
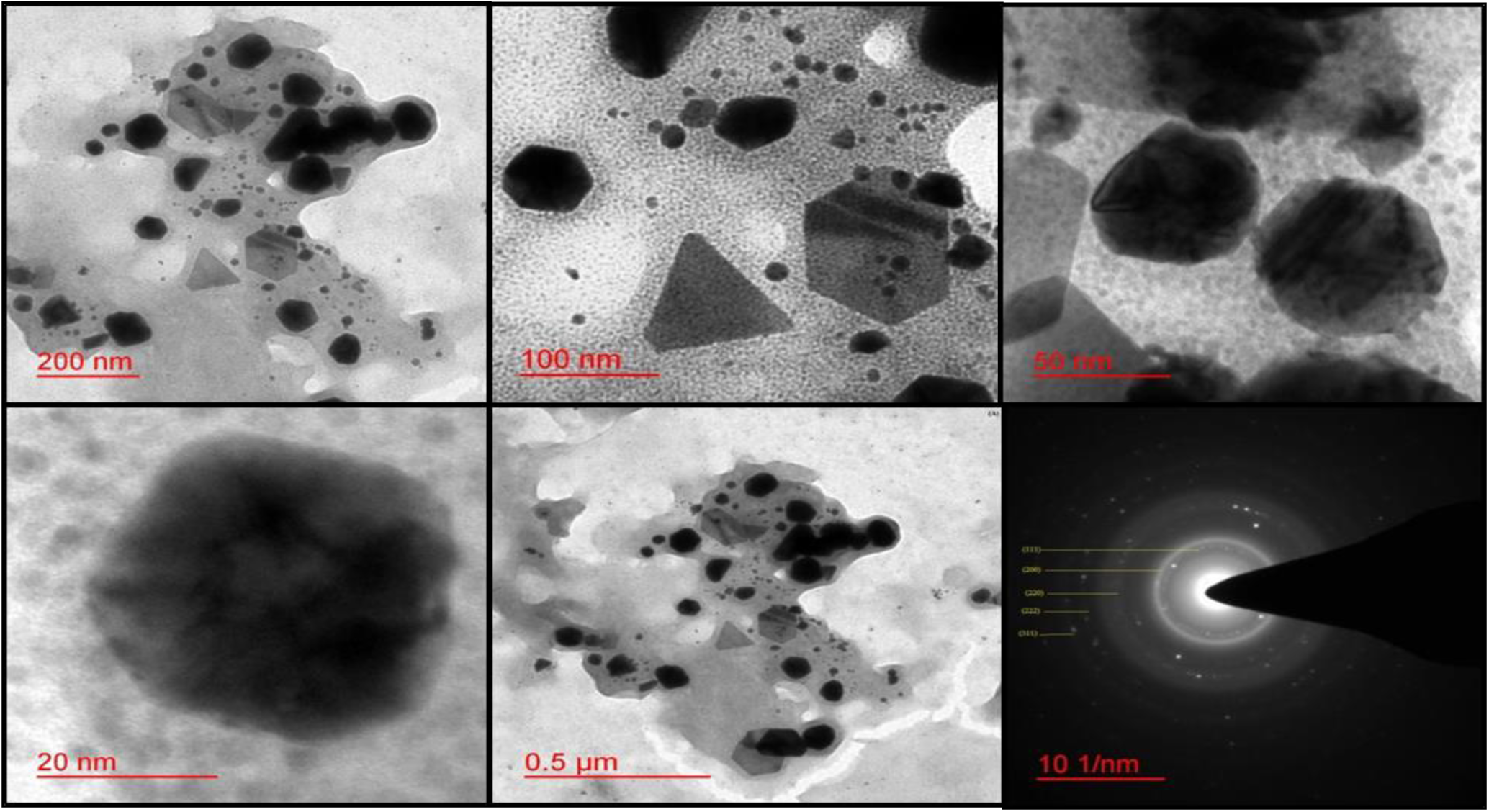
TEM morphology image of AuNPs at 200nm, 100nm, 50nm, 20nm, 0.5nm and SEAD pattern.

#### 3.3.7. SEM

SEM is used for the further characterization to determine the structural morphology, shape and size of the synthesized AuNPs (**40**). The structural morphology of *W. chinensis*-capped AuNPs is shown in **Fig. 9**. It showed the cylindrical, hexagonal, rectangular and spherical shape of the particles with various distributions of size.

**Fig. 9.**
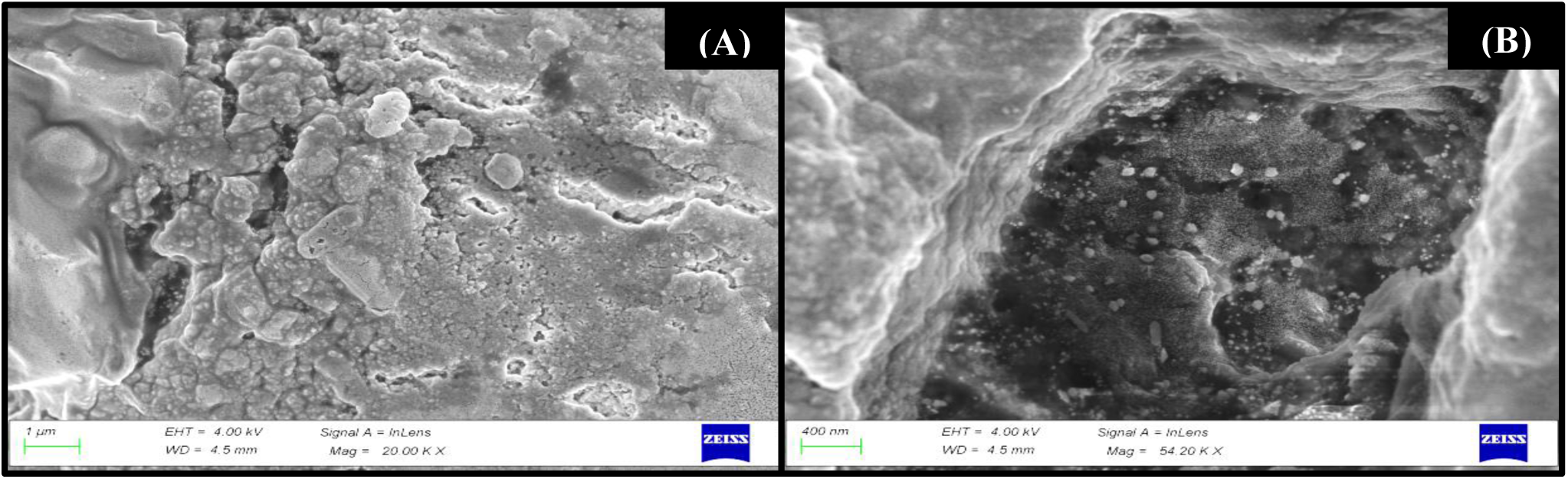
SEM images of formed AuNPs.

#### 3.3.8. X-ray diffraction analysis

The nature of the formed gold nanoparticles as a crystalline was determined by XRD analysis. The distinctive XRD pattern of AuNPs is depicted in **Fig. 10**, showing the crystalline nature of the synthesized AuNPs. From the XRD pattern, there are four distinct diffraction peaks obtained at 2θ = 38.53° (111), 43.38° (200), 65.72° (220) and 77.41° (311) Bragg’s reflection represents the planes of crystalline FCC gold metallic.

**Fig. 10.**
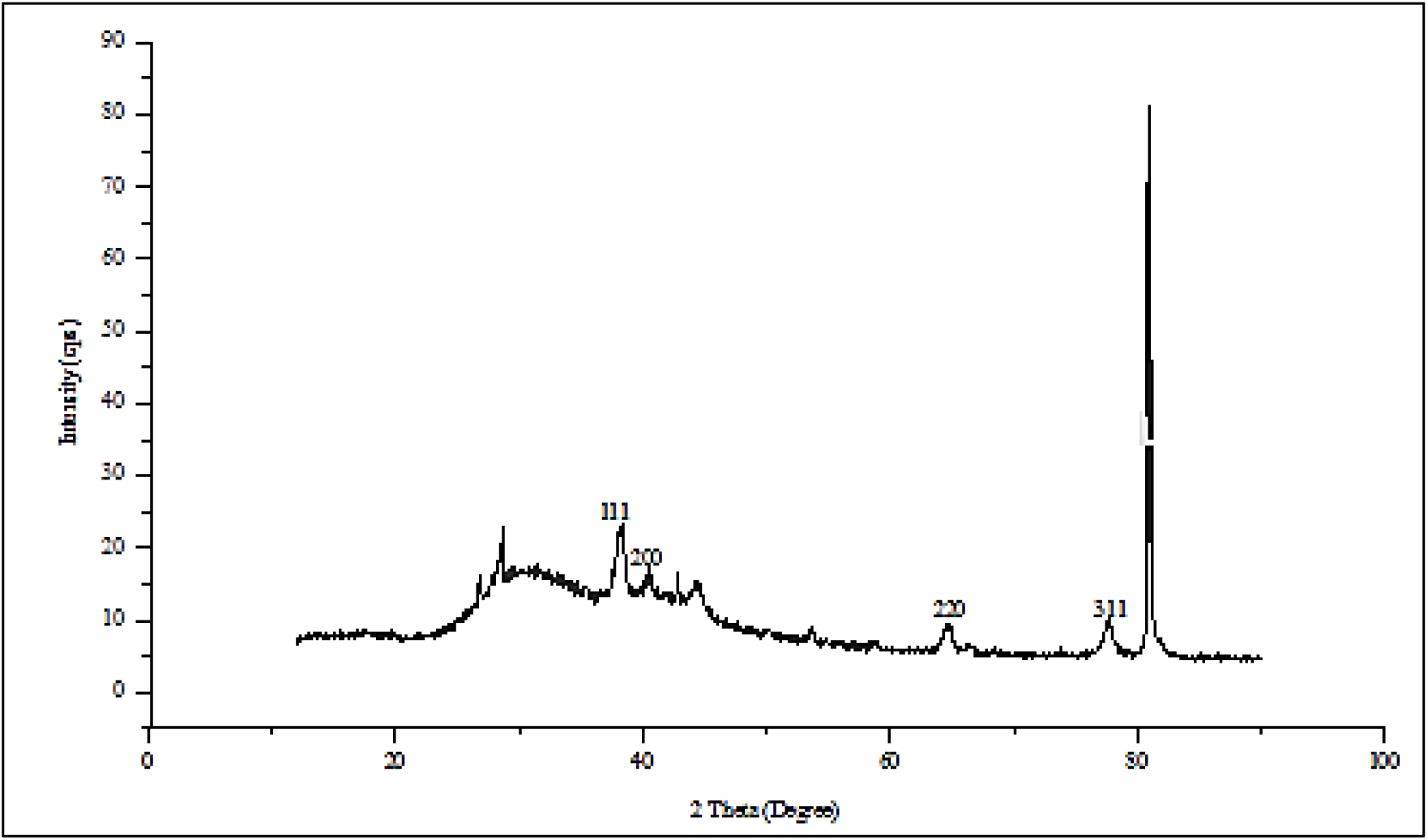
XRD pattern of gold synthesized AuNPs from internode of *W. chinensis*.

### 3.4. Antioxidant activity of the synthesized Gold nanoparticles

#### 3.4.1. DPPH activity

The scavenging assay of DPPH along with AuNPs exhibited effective inhibition activity when compared with the standard (BHT) (**Fig. 11**). Internodal extract showed greater antioxidant activity (73%) than AuNP (69%) and less than BHT taken as positive control (97%). Similarly, biosynthesized AuNPs showed less antioxidant activity than the extract (**41–35**). Lower antioxidant activity of nanoparticles may be due the participation of some metabolites of the extract as reducing and stabilizing agent.

**Fig. 11.**
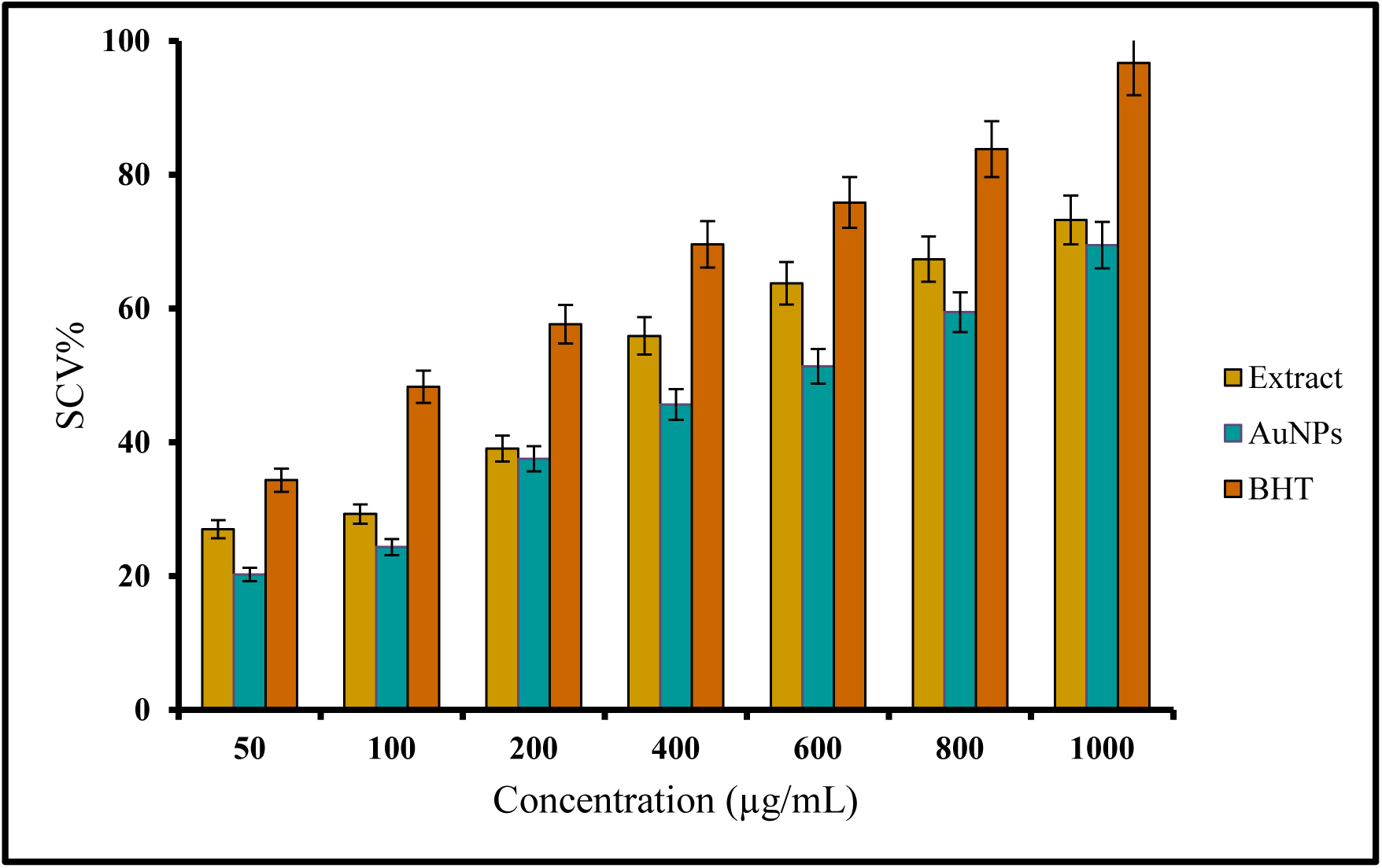
Comparison of percent inhibition of biosynthesized Internodal plant extract, AuNPs and BHT.

#### 3.4.2. Estimation of TPC and TFC Folin-Ciocalteu’s Reagent Assay and AlCl_3_ Colorimetric Method

Phenolic compounds in plant internodal extract and AuNPs were studied by measuring the total phenol content and total flavonoid content, and the results were expressed in **Fig. 12**. The amount of phenol and flavonoid content were greater in the extract as compared to AuNPs. The total phenol content in highest concentration of plant internodal extract and AuNPs (1000 µg/mL) were calculated as 613.89 µg/mL and 453.7 µg/mL, respectively (**Fig. 12A**). The total flavonoid content of plant internodal extract and AuNPs were evaluated 313.91 µg/mL and 204.12 µg/mL, respectively in the highest treatment (1000 µg/mL) (**Fig. 12B**). Our present observation shows active role of phenolic and flavonoid compounds in the formation of gold nanoparticles. Phenolic compounds show dual role as reductant as well as stabilizer during AuNP synthesis (**42**). The interaction between phenolic compounds and salt also minimizes the toxicity thus making it suitable for pharmaceutical purpose (**43**).

**Fig. 12.**
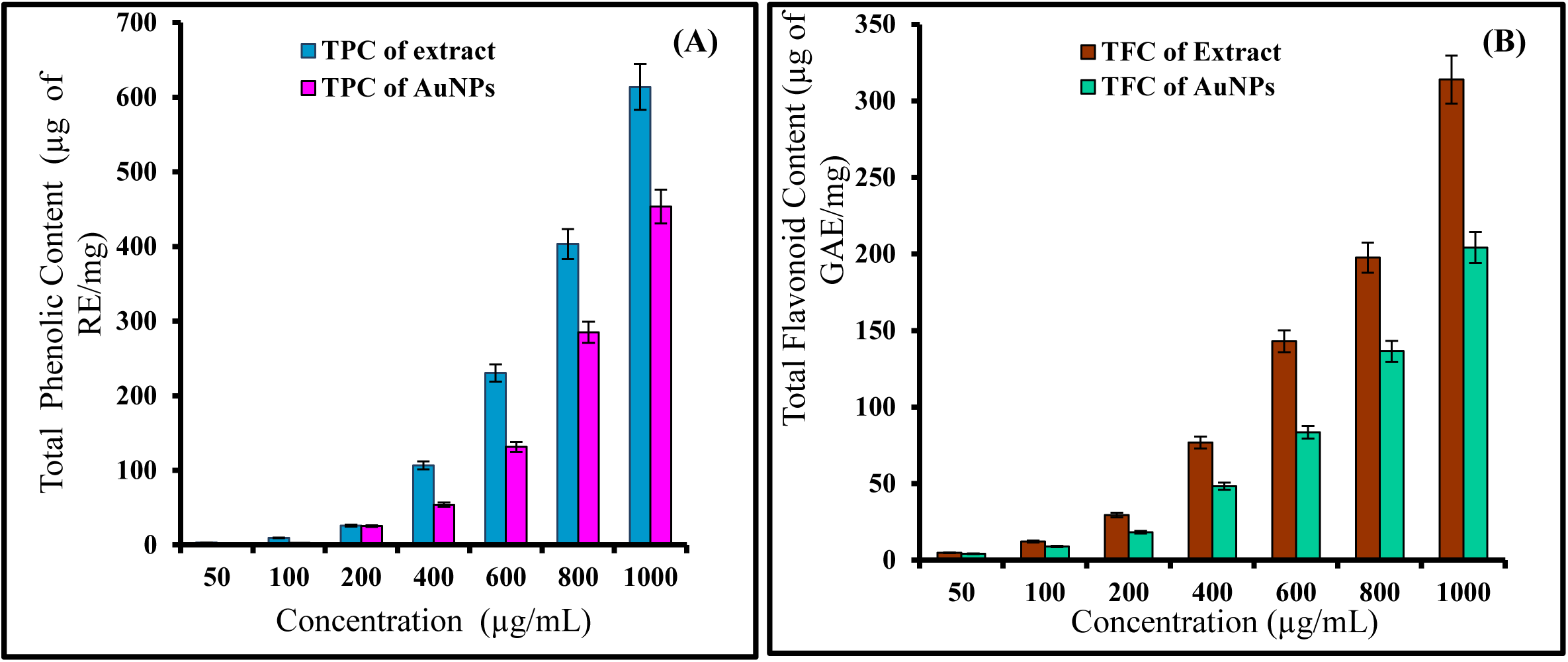
(**A**) Graph showing the percent of total phenolic content, **(B)** showing the percent of total flavonoid content.

### 3.5. Synergistic effect of Gold nanoparticles on different Bacteria strains

Antibiotics alone were not observed effective against the test organisms, but their combinations with AuNPs enhanced antibacterial activity. The combination of streptomycin with AuNPs inhibited *Klebsiella pneumoniae* and *E. coli* synergistically. The combinations of Gentamycin with AuNPs showed maximum antibacterial activity against DH5α, *Pseudomonas aeruginosa* and *Shigella boydii*. The combination of gentamycin and AuNPs acted antagonistically against MRSA (**Fig. 13a, 13b**). Enhanced antibacterial activity of antibiotic and gold nanoparticle combination will be highly helpful in handling global bacterial resistance problem (**44**). It has been observed that nanoparticles show greater antibacterial action against Gram negative strains. In present study, only MRSA is a Gram positive and combination of antibiotic and nanoparticle was not effective against it.

**Fig. 13a.**
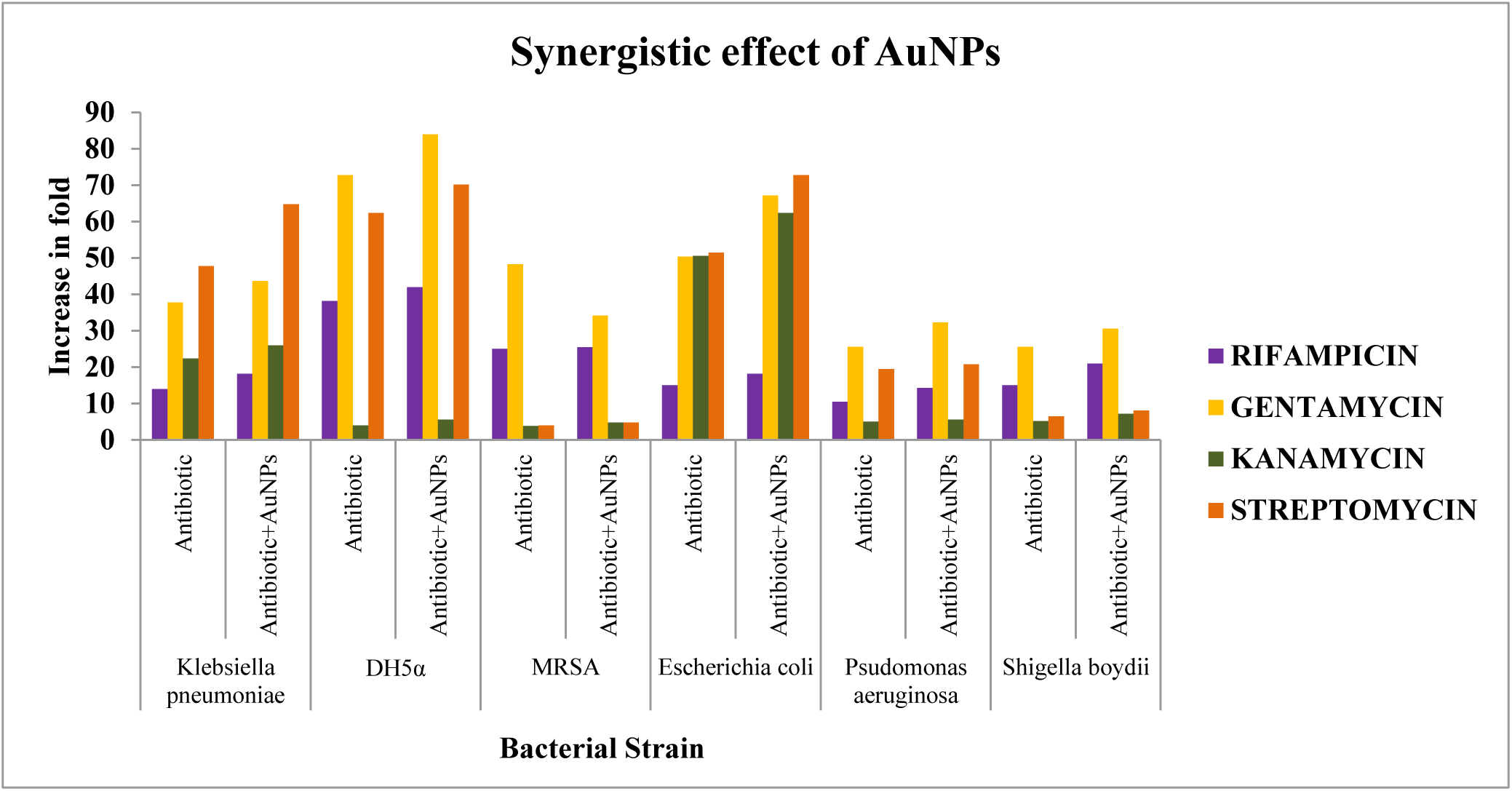
Showing synergistic effect of AuNPs against Bacterial strains

**Fig. 13b.**
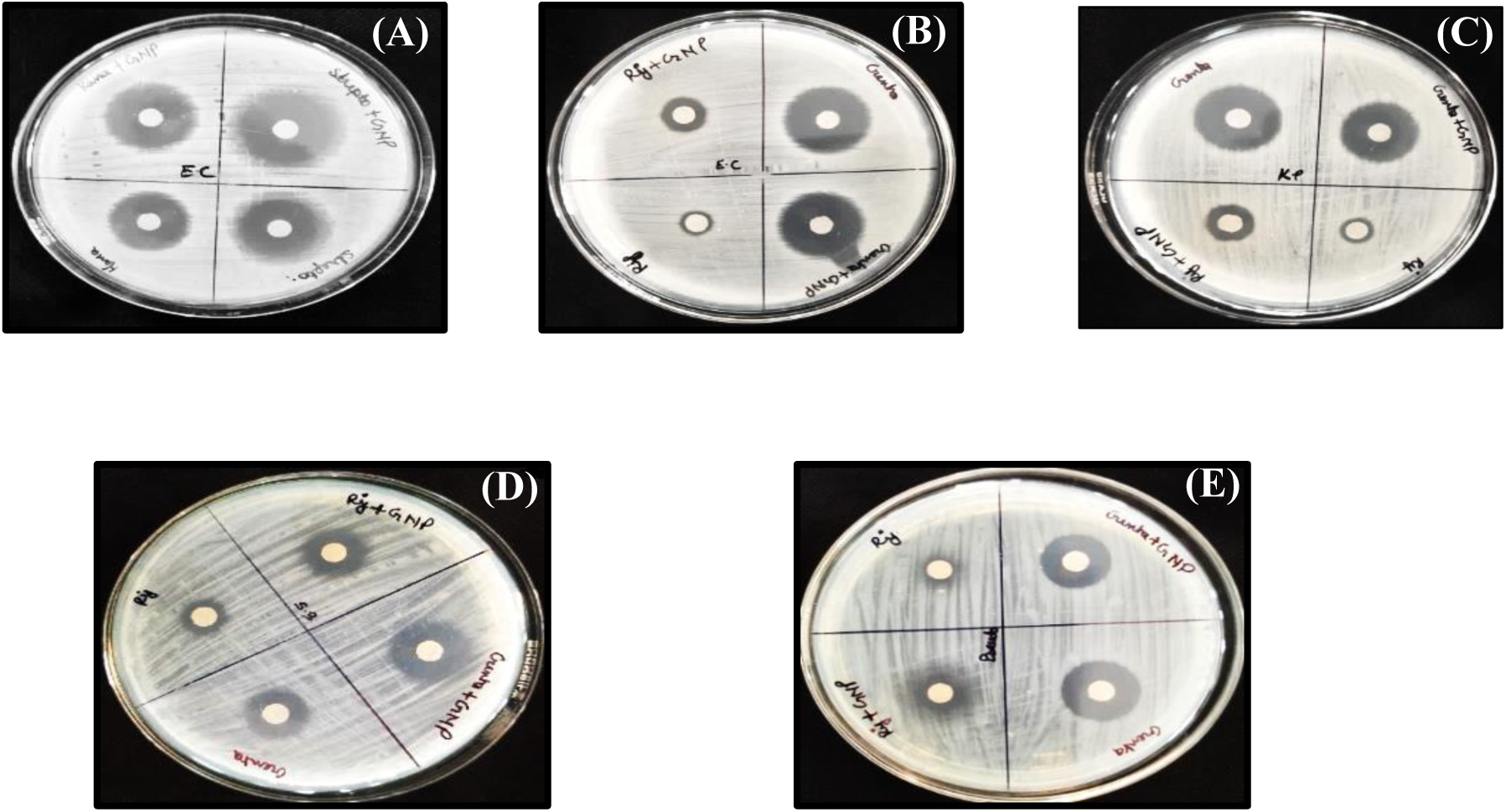
Antibacterial activity of biosynthesized AuNPs. **A**: *Escherichia coli* (EC), **B**: *Escherichia coli* (EC), **C**: *Klebsiella pneumoniae* (KP), **D**: *Shigella boydii* (SB), **E**: *Pseudomonas aeruginosa* (P).

### 3.6. Hemolytic assay of Internodal extract and gold nanoparticles

Hemolytic activity of internodal extract of *Wedelia chinensis* and salt negligible at 10 μg/ml, but hemolytic effect increases with the increase in their concentration. Gold nanoparticle (NP) showed adverse effect on RBCs and showed concentration dependent increase in 87% hemolysis (**Fig. 14**). Due to small size, free nanoparticles can enter the blood stream by penetrating alveolar lining. The interaction between NP and RBC may change cell morphology, disrupt cell membrane thus leading to hemolysis (**45**). Many researchers found that protein coating of nanoparticles causes reduction in hemolysis (**46–47**). Triton-X, taken as positive control showed 100% hemolysis.

**Fig. 14.**
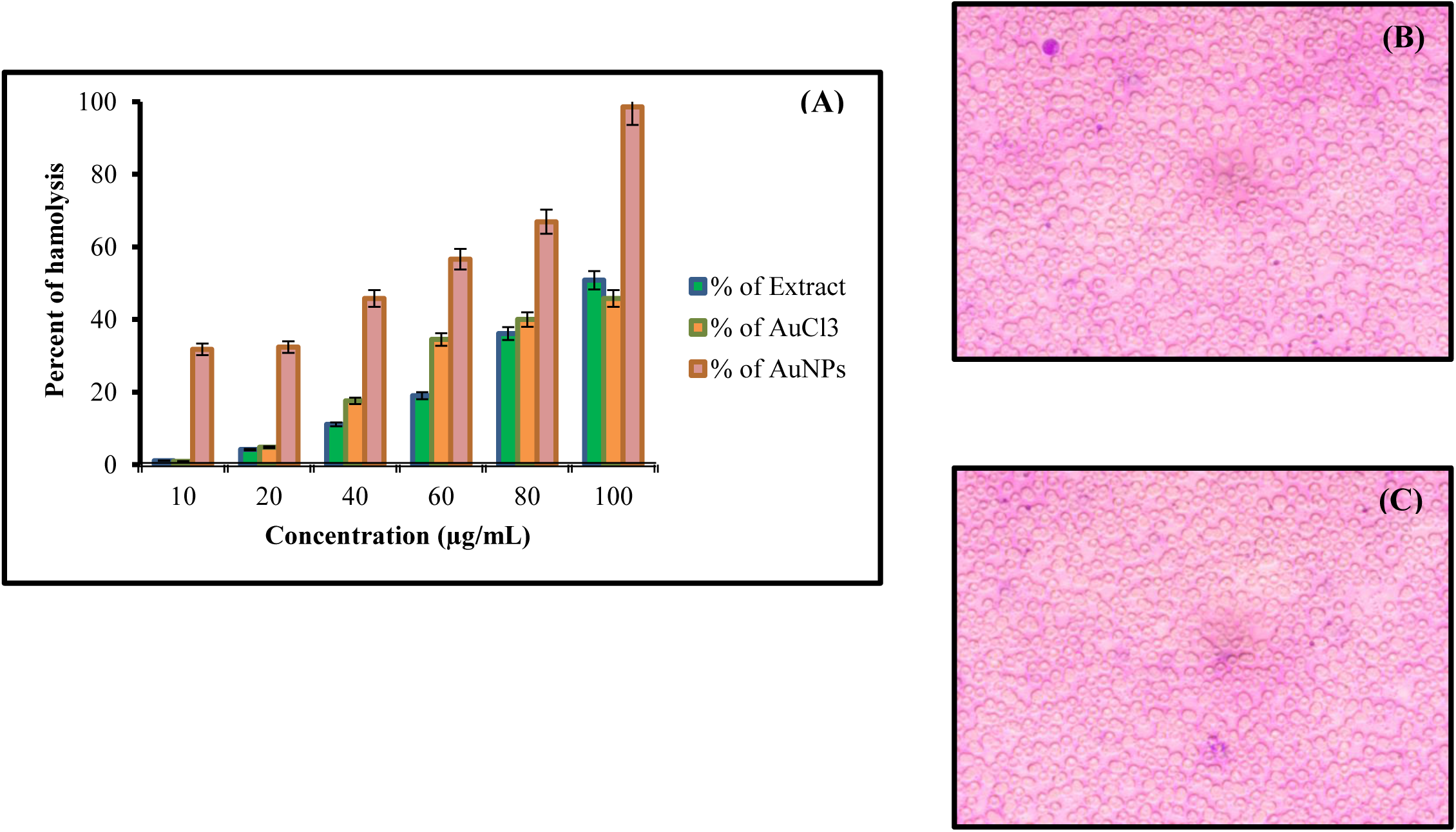
Showing **(A)** Hemolytic activity of Extract, AuCl_3_ and AuNPs; **(B)** Action of Internodal plant extract on RBC cells; **(C)** Action of AuNPs on RBC cells.

### 3.7. AuNPs Catalyzed Degradation of Methylene Blue Dye

Biogenic AuNPs prepared from *Wedelia chinensis* internode in the presence of NaBH_4_ acts as bioreducing agent for the degradation of methylene blue (MB). Discoloration and degradation of the Methylene blue dye started soon after we mixed MB with AuNPs and NaBH_4_ (**Fig. 15A**), indicating catalytic efficiency of the AuNPs. To understand the kinetics, the degradation of methylene blue (MB) was observed through UV–Vis spectrophotometer at 630 nm of wavelength at different intervals of time. **Fig. 15C**. shows the reduction of methylene blue by the gold nanoparticles through UV-vis spectrophotometer. The degradation of Methylene blue was 20 to 30% after 30 minutes. Further decrease in the intensity of absorption band at 630 nm was seen where more than 90% reduction was occurred after 3 hrs (**Fig. 15B**). The reaction follows the linear relationship as follows y =0.3325X+29.843; R^2^ =0.9911. On the basis of R^2^ value, the degradation of methylene blue by AuNPs followed second-order kinetics (R^2^=0.9911) (**Fig. 15D**). Thus, it is evident that gold nanoparticle possesses catalytic activity that reduces methylene blue at low concentration (1 mg). Gold nanoparticles are acting as absorbents of dye and they are effective in the removal of several dyes like methylene blue, methylene orange, Congo red and others (**48**). Dyes released by textile industries and other sources pollute our water bodies as organic contaminants and nanoparticles show a great application in the removal of these dyes (**49**).

**Fig. 15.**
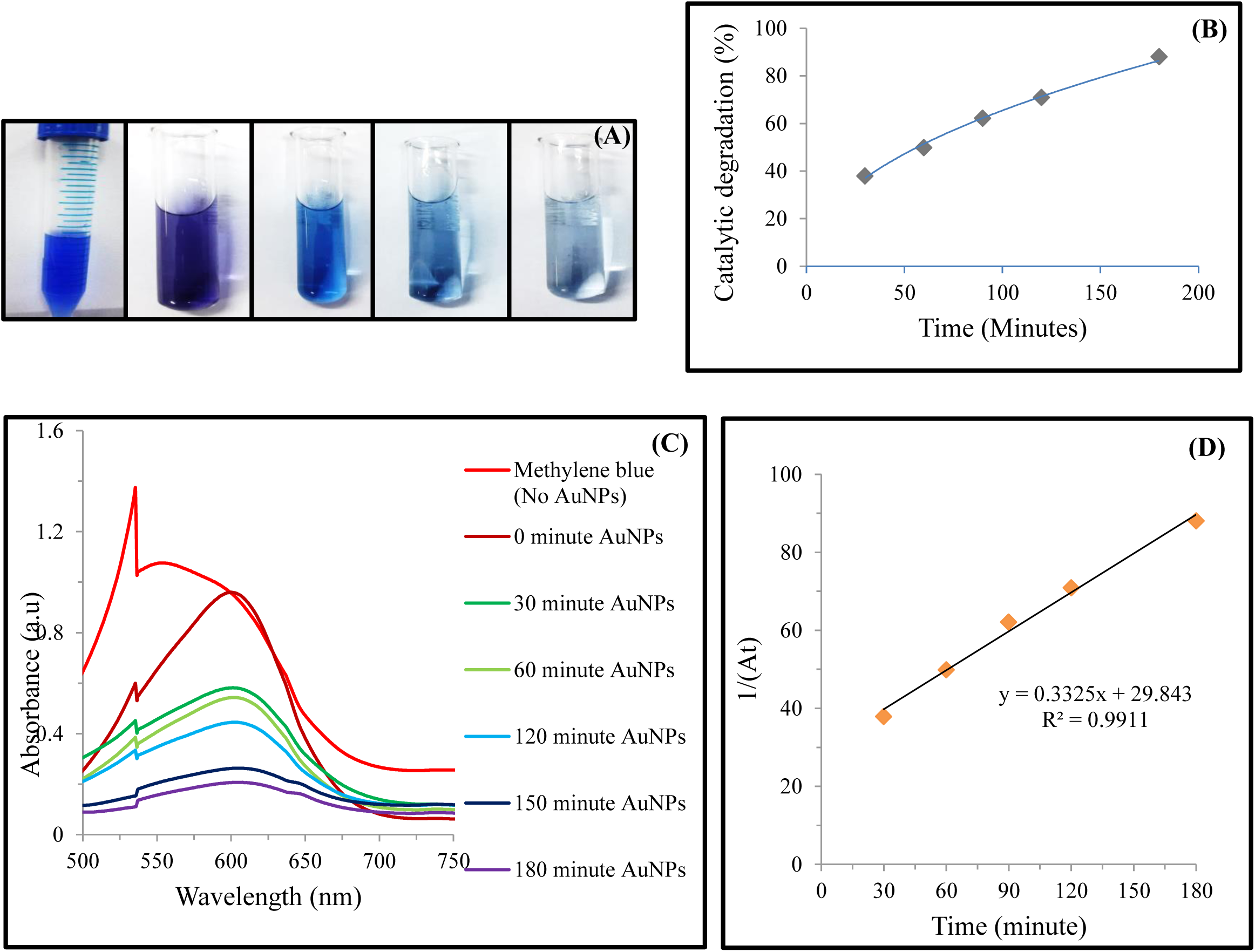
(**A**) Color change from blue to colorless (Methylene blue, AuNPs+ MB at 0 minute, AuNPs+ MB at 30 minute, AuNPs+ MB at 60 minute, AuNPs+ MB at 150 minute, AuNPs+ MB at 180 minute) showing degradation of MB dye, **(B)** Degradation of MB with time, **(C)** UV–vis absorption spectra for the reduction of MB by AuNPs at different time intervals, **(D)** Plot of 1/(At) against time for the catalytic degradation of MB by AuNPs.

## 4. Conclusion

Nanoparticles involving green technique is ecologically benign for controlling the size and form of the NPs. Biogenic interactions exhibited stability, intercalative, and antioxidant properties that had outstanding biocompatibility and bioavailability with physiologically active organic molecules. Internodal extract of *Wedelia chinensis* involves the cost effective synthesis of gold nanoparticles. The AuNPs show remarkable activity against Methylene blue dye indicates its catalytic efficiency against toxic substances. The spectrum of AuNPs toxicity suggests appropriate use in a variety of medical specialties in the future. It may also be used for anticancer, gene, or drug delivery agents since the foundation of the *W. chinensis* manufactured AuNPs demonstrated significant results. Significant synergistic antibacterial activity of gold nanoparticles was seen in combination with antibiotics. This combinational therapy will be more effective and the chances of developing the resistant bacteria against them will be minimum.

